# Short- and long-term high-fat diet exposure differentially alters phasic and tonic GABAergic signaling onto lateral orbitofrontal pyramidal neurons

**DOI:** 10.1101/2023.05.08.539875

**Authors:** L.T Seabrook, C Peterson, D Noble, M Sobey, T Tayyab, T Kenney, A.K Judge, M Armstrong, S Lin, S.L. Borgland

## Abstract

The chronic consumption of caloric dense high-fat foods is a major contributor to increased body weight, obesity, and other chronic health conditions. The orbitofrontal cortex (OFC) is critical in guiding decisions about food intake and is altered with diet-induced obesity. Obese rodents have altered morphological and synaptic electrophysiological properties in the lateral orbitofrontal cortex (lOFC). Yet the time course by which exposure to a high-fat diet (HFD) induces these changes is poorly understood. Here male mice are exposed to either short- (7 day) or long-term (90 day) HFD. Long-term HFD exposure increases body weight, and glucose signaling compared to short-term HFD or a standard chow control diet (SCD). Both short and long-term HFD exposure increased the excitability of lOFC pyramidal neurons. However, phasic and tonic GABAergic signalling was differentially altered depending on HFD exposure length, such that tonic GABAergic signaling was decreased with early exposure to the HFD and phasic signaling was changed with long-term diet exposure. Furthermore, alterations in the short-term diet exposure were transient, as removal of the diet restored electrophysiological characteristics similar to mice fed SCD whereas long-term HFD electrophysiological changes were persistent and remained after HFD removal. Finally, we demonstrate that changes in reward devaluation occur early with diet exposure. Together, these results suggest that the duration of HFD exposure differentially alters lOFC function and provides mechanistic insights into the susceptibility of the OFC to impairments in outcome devaluation.

**Significant statement:** This study provides mechanistic insight on the impact of short- and long-term high fat diet (HFD) exposure on GABAergic function in the lateral orbitofrontal cortex (lOFC), a region known to guide decision making. We find short-term HFD exposure induces transient changes in firing and tonic GABA action on lOFC pyramidal neurons, whereas long-term HFD induces obesity and has lasting changes on firing, tonic GABA and inhibitory synaptic transmission onto lOFC neurons. Given that GABAergic signaling in the lOFC can influence decision making around food, these results have important implications in present society as palatable energy dense foods are abundantly available.

## Introduction

In our modern food environment, easily accessible calorically dense foods are often high in sugars and fats (Krebs, 2009). These hyperpalatable foods are often paired with incentivizing food cues and are frequently consumed beyond metabolic need (Kanoski and Boutelle, 2022). Over time, chronic consumption of energy dense food leads to increased bodyweight and concurrent metabolic dysfunction, such as type 2 diabetes. While individual differences in genetic and environmental factors contribute to the development and maintenance of diet-induced obesity(Loos and Yeo, 2022), the primary contributor to increased body weight is the overconsumption of energy-dense foods (Swinburn et al., 2009). In our current food environment where high-caloric foods are highly palatable and easily accessible, dietary self-regulation is increasingly crucial to help influence food choices. Functions of the medial prefrontal cortex and the orbitofrontal cortex (OFC) have been implicated in various neurocognitive phenotypes, such as impulse control, decision making, and reward valuation(Knudsen and Wallis, 2022), and poorer performance on these neurocognitive tasks is associated with obesity and increased food intake (Vainik et al., 2013; Lowe et al., 2019; Seabrook et al., 2023).

The OFC plays a critical role in dietary regulation with changing internal states as it integrates reciprocal inputs and outputs from mesolimbic, sensory, and striatal regions (Izquierdo, 2017). OFC neurons respond to satiety status, with an increase in hunger-related activity in rats (de Araujo et al., 2006), monkeys (Critchley and Rolls, 1996), and humans (O’Doherty et al., 2000). Furthermore, neurons in the OFC initially respond to food rewards (Schoenbaum et al., 1998) and after learning, fire in response to anticipatory events (de Araujo et al., 2006). Mice increase sucrose licking responses when OFC neurons that previously responded to caloric rewards were activated (Jennings et al., 2019), suggesting that OFC neuronal activity is sensitive to the value of food and update actions to obtain food.

The lateral orbitofrontal cortex (lOFC) is comprised of layered pyramidal neurons with their excitability being modulated by local GABAergic interneurons and astrocytes(Quirk et al., 2009; Lau et al., 2021). In the obese state, lOFC pyramidal neurons have altered structural (Thompson et al., 2017) and synaptic properties (Thompson et al., 2017; Lau et al., 2021; Seabrook et al., 2023). There is a decrease in inhibitory presynaptic release probability onto pyramidal neurons resulting from altered astrocytic glutamate GLT-1 transporter function leading to increased endocannabinoid tone via mGluR5 activation (Lau et al., 2021). Furthermore, obesity induces a decrease in tonic GABA resulting in disinhibited pyramidal neurons (Seabrook et al., 2023). Impaired synaptic function was restored by increasing tonic GABA (Seabrook et al., 2023) or by increasing astrocyte GLT-1 function (Lau et al., 2021). While these findings demonstrate a synaptic mechanism on how the lOFC is altered in obesity (Thompson et al., 2017; Lau et al., 2021; Seabrook et al., 2023), it is unknown if these events occur early with diet exposure, or if this only occurs in the obese state. Identifying the time course of these changes may help us identify ways to intervene earlier to prevent the behavioural sequelae associated with disinhibition of the OFC (Bissonette et al., 2015; Seabrook et al., 2023). Therefore, using patch clamp electrophysiology in lOFC brain slices from mice fed a 7- or 90-day high-fat diet (HFD) or their age matched controls, we determined the time course of neurophysiological and behavioral changes in the development of obesity.

## Materials and Methods

### Animals

All protocols were in accordance with the ethical guidelines established by the Canadian Council for Animal Care and were approved by the University of Calgary Animal Care Committee. Adult male C57BL6 mice were obtained from Charles Rivers Laboratories or from the Clara Christie Centre for Mouse Genomics (University of Calgary). Mice were maintained on a 12 h light/dark schedule (lights on at 8:00 A.M. Mountain Standard Time). All experiments were performed during the light cycle of the animals. Animals were typically group housed (2-5 animals per cage). However, because a staggered design disrupted the cage social structure and resulted in fighting, mice were singly housed during last 7 days (90 day HFD, or SCD) or the entire 7-day diet manipulation, to allow for use of a single mouse per day for electrophysiology experiments.

### Diets

The control diet was obtained from Lab diets (Somerville, NJ, Rodent Diet #5062) composed of 23% protein, 55 % carbohydrates, and 22% fat (3.76kcal/g). The HFD was obtained from Research diets (New Brunswick, NJ). The HFD (D12492) was composed of 20% protein, 20% carbohydrate, and 60% fat (5.21kcal/g). Mice given 90 days of HFD were given the diet P60-P150, whereas mice given the diet for 7 days were delivered the diet P143-P150. In the conditioned taste aversion experiment, mice were given 7-day low fat diet (LFD, Research diets D12450J: composed of 20% protein, 70% carbohydrate and 10% fat from calories (3.82kcal/g)) instead of SCD. In some experiments, after 7- or 90-day HFD exposure, mice were returned to a standard chow diet for 7 days before electrophysiological recordings.

### Glucose tolerance test

In a separate cohort, mice were exposed to standard chow only, 7-day or 90-day HFD were fasted overnight. A baseline blood sample was collected from the tail vein (a small cut 1-2mm from the end of the tail, time 0). Mice were then administered an intraperitoneal injection 20% glucose solution (20% D-glucose in 0.09% saline, 2g of glucose per kilogram of body weight) and blood was collected at 15, 30, 45, 60, 75, 90, 105, 120, 150, 180, 210 minutes post-injection). Blood glucose levels were measured with an Accu-Chek Aviva blood glucose meter.

### Electrophysiology

All electrophysiology recordings were performed in slice preparations containing the lOFC from 5-6 month old mice fed 0, 7- or 90-days HFD. Mice were anesthetised with isoflurane and transcardially perfused with an ice-cold N-methyl-D-glucamine (NMDG) solution of the following composition (in mM): 93 NMDG, 2.5 KCl, 1.2 NaH2PO4.H2O, 30 NaHCO3, 20 HEPES, 25 D-glucose, 5 sodium ascorbate, 3 sodium pyruvate, 2 thiourea, 10 MgSO4.7H2O, and 0.5 CaCl2.2H2O. Mice were quickly decapitated, brains were extracted and 250 µm coronal sections containing the lOFC were prepared using a vibratome (Leica, Nussloch, Germany) in the same ice cold NMDG solution. Slices recovered in warm NMDG solution (32°C) for 10min, before being transferred to a long-term holding chamber containing artificial CSF (ACSF) of the following composition (in mM): 126 NaCl, 1.6 KCl, 1.1 NaH2PO4, 1.4 MgCl2, 2.4 CaCl2, 26 NaHCO3, and 11 glucose (32–34°C). All solutions were saturated with 95% O2/5% CO2. Before recording, sections were then transferred to the recording chamber and super fused with aCSF maintained at 32°C. lOFC cells were visualized on an upright microscope (model BX51WI-Olympus) using “Dodt-type” gradient contrast infrared optics and whole-cell recordings were made using a MultiClamp 700B amplifier (Axon Instruments, Union City, CA) and collected with pClamp10. Pyramidal neurons in layer II/III were identified by morphological characteristics of a large soma size and triangular shaped appearance, as well as electrophysiological properties of high capacitance (>100 pF) and were recorded approximately 100-300 µm above the inflection point of the rhinal sulcus.

For current clamp experiments (cellular activity), recording electrodes (3-5 MΩ) were filled with (in mM) 130 potassium-D-gluconate, 10 KCl, 10 HEPES, 0.5 EGTA, 10 sodium creatine phosphate, 4 Mg-ATP and 0.3 Na2GTP. After breaking into the cell, membrane resistance was recorded in voltage clamp. Cells were then switched into current clamp mode and the resting membrane potential was recorded. The resting membrane potential as defined as the potential generated across the cell membrane by the difference in charge from internal and external solutions was calculated 2 minutes after achieving whole cell configuration by the amplifier. Membrane potential for each neuron was set to -70 mV by current injection via the patch amplifier. A current step protocol consisting of 21 steps (0-500pA, 25pA increments, 400ms in duration, 3 seconds apart) was applied and the number of action potentials at each step was recorded. For some experiments, the current step protocol was initiated after application of picrotoxin (100 μM).

For voltage clamp experiments including measurements of spontaneous inhibitory postsynaptic currents (sIPSCs), miniature inhibitory postsynaptic currents (mIPSCs) and tonic GABA currents, recording electrodes were filled with a cesium chloride (CsCl) internal solution consisting of (in mM) 140 CsCl, 10 HEPES, 0.2 EGTA, 1 MgCl_2_, 2 MgATP, 0.3 NaGTP, 5 QX-314-Cl. sIPSC currents were recorded at -70 mV in the presence of DNQX (10 μM) to block AMPA receptors, strychnine (1 μM) to block glycine receptors, DPCPX (1 μM) to block adenosine A1 receptors, and CGP-35348 (1 μM) to block GABA_B_ receptors. mIPSCs were also recorded in the presence of tetrodoxin (TTX; 500 nM) to block action potential firing. IPSCs were filtered at 2 kHz, digitized at 10 kHz and collected on-line using pCLAMP 10 software. For tonic GABA experiments gabazine (100 μM) was added to aCSF bath solution which included DNQX (10 μM), strychnine (1 μM), DPCPX (1 μM), CGP-35348 (1 μM) and APV (10 μM). The change in holding potential and RMS noise was obtained by a Gaussian fit to an all points histogram over a 5 second interval(Herman and Roberto, 2016). In some experiments, 4,5,6,7-tetrahydroisoxazolo[5,4-c]pyridin-3-ol (THIP; 5 μM) was washed onto the slice. The junction potential of +4 mV for CsCl internal or +16.2 mV for KGluconate internal solution was not corrected. Recordings exhibiting a >20% change in series resistance were discarded. GABA_A_ sIPSCs were quantified from tonic GABA recordings (pre-gabazine) and were selected for amplitude (>12 pA), rise time (<4 ms), and decay time (<10 ms) using the MiniAnalysis program (Synaptosoft). mEPSCs were selected for amplitude (>12pA), rise time (<4ms) and decay time (<6 ms) using MiniAnalysis Mini60 program (Synaptosoft).

### Devaluation by conditioned taste avoidance

To test for conditioned taste avoidance (CTA), lean adult C57BL/6 mice underwent a 6-day taste avoidance-conditioning paradigm (see detailed methods: Seabrook et al 2023). Briefly, three days prior to conditioning, mice were exposed to 5.15% grape and orange Kool-Aid^TM^ flavoured gelatine (Knox Gelatine) in their home cages to reduce food neophobia. To minimize stress, animals were brought to the testing room and remained in the room for 1 h before conditioning. During conditioning, each animal consumed a flavour of gelatine that was either paired with sickness inducing lithium chloride (LiCl) (40mL/kg of 0.24M LiCl, i.p.) or vehicle (40mL/kg 0.9% saline, i.p) over 3 conditioning days. Each animal received the paired injection with one flavour and the unpaired injection with a different flavour every second day for a total of 6 consecutive days of injections. Flavours paired with LiCl were counterbalanced. The two different gelatine exposures were administered in distinct environmental contexts. One context consisted of a smooth cage bottom, a paper house, and gelatine placed in a square plastic weigh boat. The second context had white paper towel on the cage bottom and gelatine was delivered in a plastic circle weigh boat. After 1h access to the flavoured gelatine, the remaining food was removed and weighed, and mice were immediately injected with either vehicle (VEH) or LiCl. Mice were then placed back into their conditioning cage for 1h before being moved to their home cage. Test days were counterbalanced and consisted of exposure to either orange or grape flavour on day 1 and orange or grape flavour on day 2. In the diet experiments, after CTA, mice were randomized to either a HFD or a LFD for 7 days. Mice were then re-exposed to grape or orange gelatine for 1 h on separate days and gelatine consumption was measured. The “valued” state was exposure to the unpaired gelatine flavour, whereas the “devalued” state was exposure to the LiCl-paired gelatine flavour.

### Devaluation by satiety

Mice were mildly food restricted and maintained at 85% of their original weight throughout training and testing. Instrumental responding for a 30% sucrose solution in was performed in sound- and light-attenuated Med-associate chambers (St-Albans, Vermont) equipped with a retractable active lever. A cue light was positioned above the lever and was illuminated when the lever was active. Chambers were illuminated with a house light, which signalled the beginning of the session. With the appropriate number of lever presses, a 30% liquid sucrose (dissolved in H_2_O) was delivered in 0.1 ml increments into the cup via a syringe connected to a pump. All training consisted of 1h sessions. To train animals to lever press, we shaped behaviour by baiting the lever with sucrose during a fixed ratio (FR) 1 schedule of reinforcement, whereby 1 lever press delivered 1 liquid sucrose outcome. To escalate responding, we switched to a random ratio (RR) 5, 10, and 20 schedules of reinforcement. During RR, outcomes are delivered, on average, every 5, 10, or 20 lever presses, but not precisely every 5, 10, or 20 times the lever is pressed. Devaluation by satiety occurred over two testing days and the “valued” and “devalued” conditions were counterbalanced for every diet and manipulation. Water was removed prior to behavioural testing (zeitgeber time (ZT) 9). At ZT 21, a bottle of either H_2_0 (“valued”) or 30% sucrose (in H_2_0, “devalued”) was introduced into the cage and the animals were allowed to drink freely for 3 hours. Further, if there were 0 or 1 lever presses in the valued condition, mice were excluded due to a lack of engagement in the task. One 90-HFD mouse was excluded because total lever pressing over the valued and devalued days met the outlier criteria using a Grubbs outlier statistical test. Mice were given access to SCD or HFD throughout training and testing for the SCD or 90 days HFD group. Mice were given access to 7 days HFD in the days preceding and during the devaluation test.

### Data analysis and statistics

Data are expressed as the mean ± SEM. Individual data points are overlayed averages where possible. All grouped data were analyzed with a one-way ANOVA except in the glucose tolerance test, behavioural tests, and excitability F-I plots where a two-way ANOVA was employed. When appropriate, data was analyzed with a Tukey’s post hoc comparison test or a Holm’s Sidak test (behavioural tests) following the ANOVAs. All statistical analyses were performed in GraphPad Prism 9.4.1 (GraphPad, US) and presented in the figure legends. To determine the mean excitability slope, R^2^ value of individual cells determined by non-linear regression whereby x = current step and y= frequency of action potentials. All significance was set at P<0.05. Individual responses are plotted over averaged responses. Experimental designs and samples sizes were aimed at minimizing usage and distress of animals and were sufficient for detecting robust effect sizes.

## Results

### Long-term HFD exposure increases body weight and decreases glucose clearance

To examine the effects of a HFD on lOFC pyramidal neurons adult mice were fed either a HFD for 0-, 7- or 90-days (Figure 1a). Long-term HFD increases body weight and induces metabolic dysfunction (Seabrook et al., 2023). To address if short-term (7 days) or long-term (90 days) HFD exposure differentially alters blood glucose clearance, a proxy for insulin signaling, mice were given a glucose challenge and blood glucose was measured every fifteen minutes for two hours. Mice fed 90 days of a HFD had increased bodyweight (Figure 1b), as well as decreased glucose clearance after a glucose challenge (Figure 1c) and increased area under the curve (Figure 1d), indicating that long-but not short-term HFD exposure increases body weight and impairs glucose clearance.

**Figure 1:**
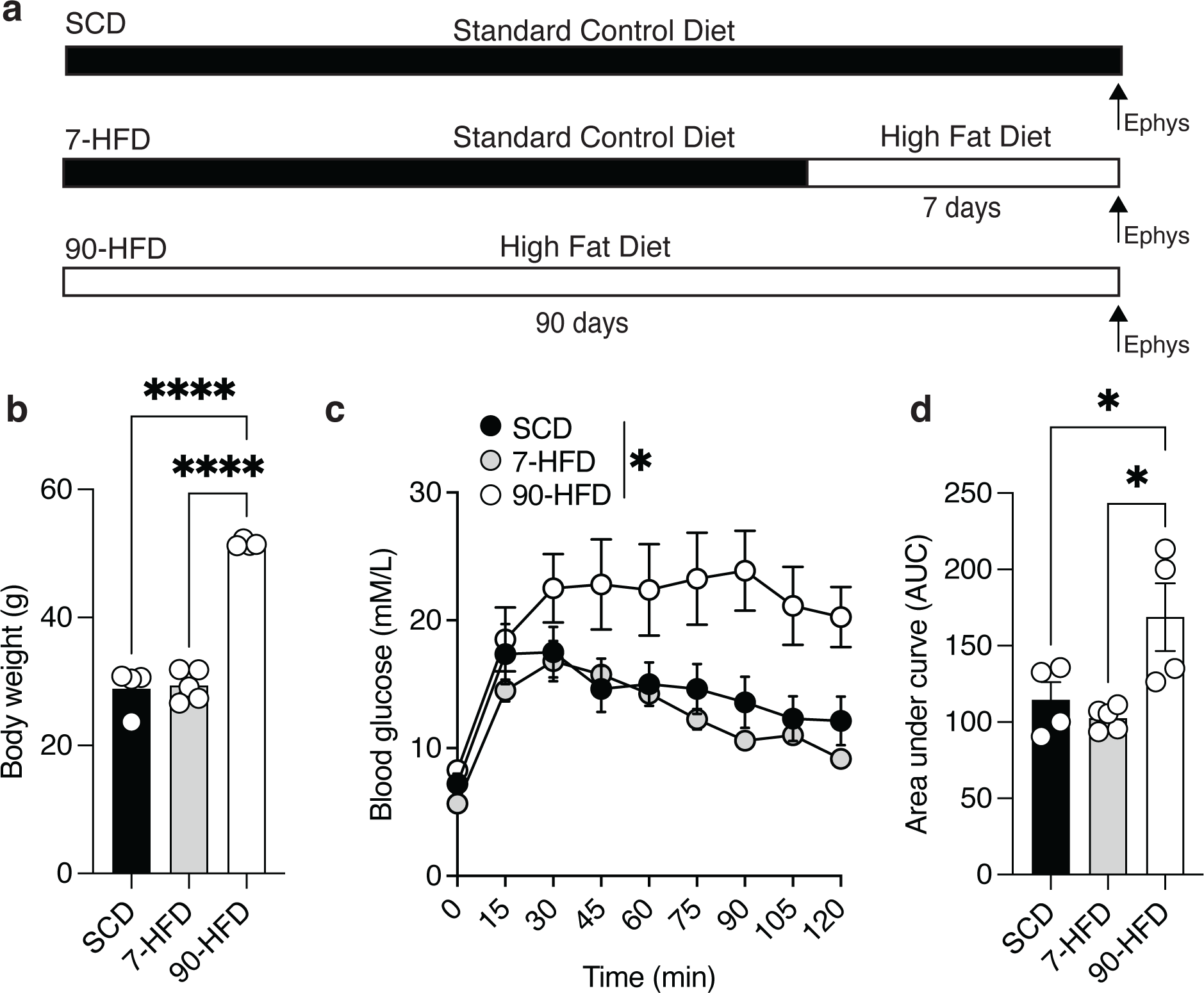
Long-term high fat-diet exposure increases body weight and decreases glucose clearance. **a)** Schematic of diet schedule. **b)** 90-day high fat diet (HFD) exposure (N=4) increased bodyweight compared to 7-day HFD (N=5) and standard chow diet (SCD) (N=4). One-way ANOVA: F (2, 10) = 115.0 P<0.0001 **** Tukey’s post hoc comparison show a difference between SCD and 90-day HFD P<0.0001 **** and 7-day HFD and 90-day HFD P<0.0001 ****. Bars represent mean and symbols represent individual values. **c)** 90-day HFD exposure (N=4) reduced glucose clearance compared to 7-day HFD (N=5), and SCD (N=4) as indicated by blood glucose (nM/L) concentrations following an intraperitoneal injection of 20% D-glucose solution. Two-way RM ANOVA: Time effect: F (3.07, 30.69) = 24.44, P<0.0001****, Diet effect F (2, 10) = 7.56, P=0.01* Time x diet interaction: F (16, 80) = 2.82, P=0.0012**. Symbols represent mean + S.E.M. **d)** 90-day HFD exposure (N=4) reduced insulin sensitivity compared to 7-day HFD (N=5), and SCD (N=4) indicated by the area under the curve (AUC). One-way ANOVA: F (2, 10) = 6.84, P=0.013 *, Tukey’s post hoc comparisons show a significant difference between SCD and 90-day HFD P= 0.049* and 7-day HFD and 90-day HFD P= 0.013*. Bars represent mean + S.E.M, symbols represent individual values.

### Short- and long-term HFD exposure increases excitability of lOFC pyramidal neurons

Our previous work demonstrated that obese mice fed a HFD have increased excitability of lOFC pyramidal neurons (Seabrook et al., 2023), but the time course of these changes in the lOFC is unknown. Current-evoked firing frequency was increased in lOFC from mice exposed to either 7 days or 90 days of a HFD compared to SCD (Figure 2a, b). Increased excitability was evident by the difference in slopes when frequency-current (F-I) plots were fitted with a non-linear regression (Figure 2c). Consumption of a HFD could alter passive membrane properties of lOFC neurons, as such we measured action potential (AP) characteristics. There was no difference in latency or threshold to fire, AP or after hyperpolarization potential (AHP) height, resting membrane potential, input resistance, or capacitance (Figure 2d-f,h,i,k,l). However, we did observe a decrease in AP and width in mice fed 90 days of a HFD (Figure 2g) and AHP width in mice fed 7 or 90 days of a HFD (Figure 2j). Furthermore, exposure to 7 or 90 days HFD decreased the rheobase, reflecting the current at which the first action potential fires (Figure 2h). Consistent with our previous findings, diet exposure increases the excitability of pyramidal neurons (Seabrook et al., 2023), but this effect occurs with diet exposure prior to obesity.

**Figure 2:**
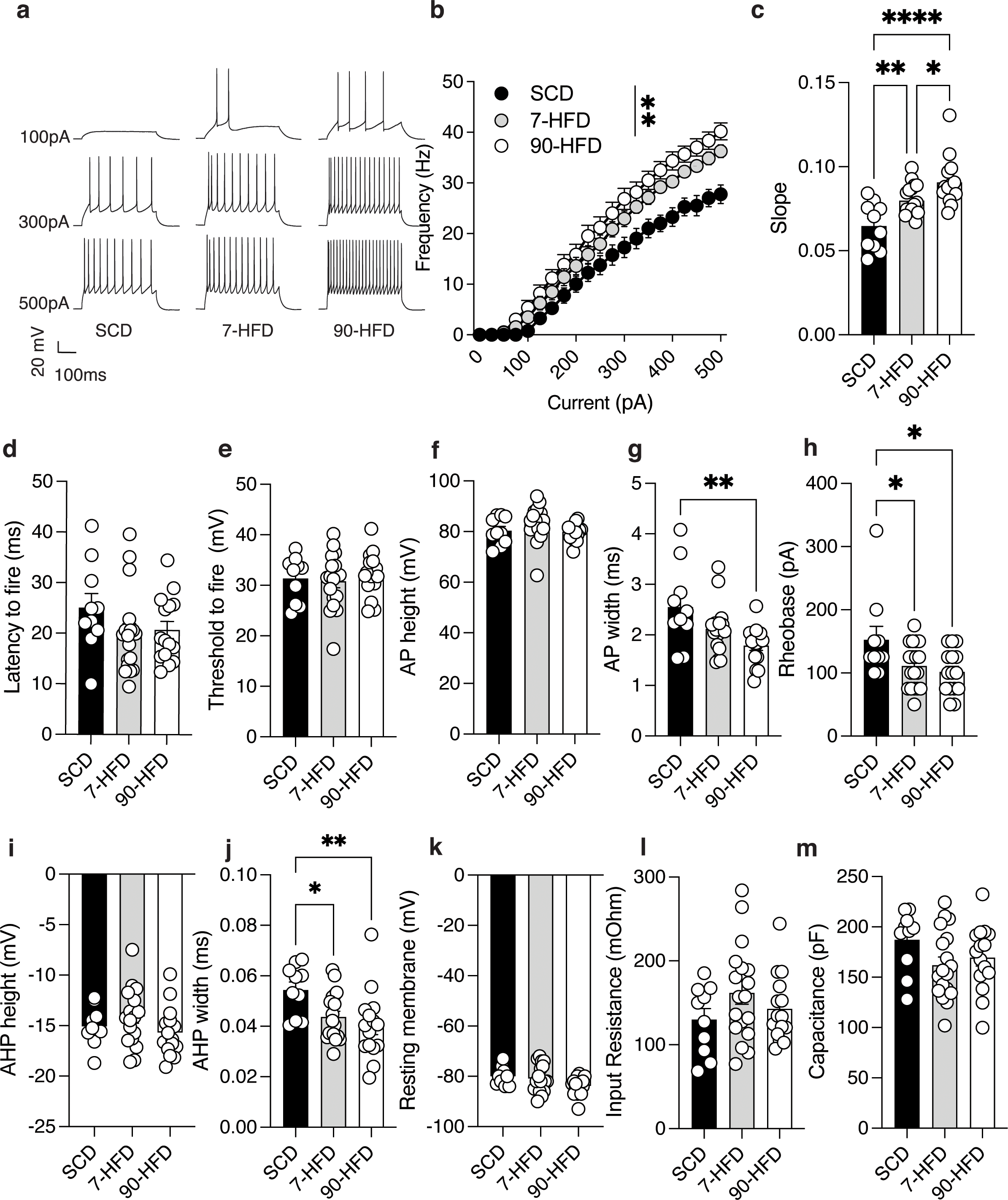
Short- or long-term high fat diet exposure increases excitability of lOFC pyramidal neurons. **a)** Representative recordings of action potentials observed at 100pA, 300pA and 500pA current steps from lOFC pyramidal neurons of SCD, 7-day HFD, and 90-day HFD. **b)** 7 days (n/N=18/7) and 90 days (n/N=15/4) of a high fat diet increased the excitability of lOFC pyramidal neurons compared to SCD (n/N=10/3) as indicated by frequency-current (F-I) plot of action potentials at current injections from 0 pA to 500pA over a 400 ms step. Two-way RM ANOVA: pA injected effect: F (2.76, 110.6) = 535.4, P<0.0001****, Diet effect: F (2, 40) = 6.71 P=0.0031 ** pA injected x diet interaction F (40, 800) = 4.07, P<0.0001****. **c)** 7 days (n/N=18/7) or 90 days (n/N=15/4) of high fat diet exposure increased the excitability of lOFC pyramidal neurons compared to SCD (n/N=10/3) indicated by difference in slope obtained from a non-linear regression fit. One-way ANOVA: F (2, 40) = 13.83, P<0.0001***. Tukey’s post hoc test showed a significant difference between SCD and 7-day HFD P=0.0078**, SCD and 90-day HFD P<0.0001**** and 7-day and 90-day HFD P = 0.038*. **d)** Diet exposure SCD (n/N=10/3), 7-day (n/N=18/7) and 90-day (n/N=15/4) HFD did not alter the latency to fire. One-way ANOVA: F (2, 40) = 1.46, P=0.25. **e)** Diet exposure SCD (n/N=10/3), 7-day (n/N=18/7) and 90-day (n/N=15/4) HFD did not alter the threshold to fire. One-way ANOVA: F (2, 40) = 0.28, P=0.75. **f)** Diet exposure SCD (n/N=10/3), 7-day (n/N=18/7) and 90-day (n/N=12/4) HFD did not alter the AP height. One-way ANOVA: F (2, 40) = 1.77, P=0.18. **g)** 90-day (n/N=15/4) HFD increased AP width compared to SCD (n/N=10/3) and 7-day HFD (n/N=18/7). One-way ANOVA: F (2, 40) = 5.94, P=0.0055. Dunnett’s post hoc comparison test showed a significant difference between SCD and 90-day HFD P= 0.0025 **. **h)** 90-day (n/N=15/4) or 7-day (n/N=18/7) HFD had decreased rheobase compared to SCD (n/N=10/3). One-way ANOVA: F (2, 40) = 4.24, P=0.021*. Dunnett’s post hoc comparison test showed a significant difference between SCD and 7-day HFD, P = 0.041*, SCD and 90-day HFD, P= 0.014*. **i)** Diet exposure SCD (n/N=10/3), 7-day (n/N=18/7) and 90-day (n/N=15/4) HFD did not alter the AHP height. One-way ANOVA: F (2, 40) = 1.65, P=0.21. **j)** 90-day (n/N=15/4) HFD or 7-day (n/N=18/7) HFD had decreased AHP width compared to SCD (n/N=10/3). One-way ANOVA: F (2, 40) = 4.93, P=0.012*. Dunnett’s post hoc comparison test showed a significant difference between SCD and 7-day HFD, P = 0.037*, SCD and 90-day HFD, P= 0.007**. **k)** Diet exposure SCD (n/N=10/3), 7-day (n/N=18/7) or 90-day (n/N=15/4) HFD did not alter the resting membrane potential. One-way ANOVA: F (2, 40) = 3.06, P=0.079. **l)** Diet exposure SCD (n/N=10/3), 7-day (n/N=18/7) or 90-day (n/N=15/4) HFD did not alter the input resistance. One-way ANOVA: F (2, 40) = 1.49, P=0.24. **m)** Capacitance was similar between SCD (n/N=10/3), 7-day (n/N=18/7) or 90-day (n/N=15/4) HFD. One-way ANOVA: F (2, 40) = 1.88, P=0.17.

In addition to changes in membrane properties, increased excitatory drive could underlie increased excitability after HFD exposure. Therefore, we quantified miniature excitatory post synaptic currents (mEPSC) onto lOFC pyramidal neurons. There was no difference in mEPSC frequency or amplitude onto pyramidal neurons from mice exposed to SCD, 7, or 90 days of HFD (Figure 3a-c). Cumulative probability plots also found no difference in inter-event interval (Figure 3d) but did find a difference in cumulative probability of amplitude (Figure 3e). These results suggest that changes in glutamatergic input is unlikely to be contributing to increased pyramidal neuron excitability.

**Figure 3:**
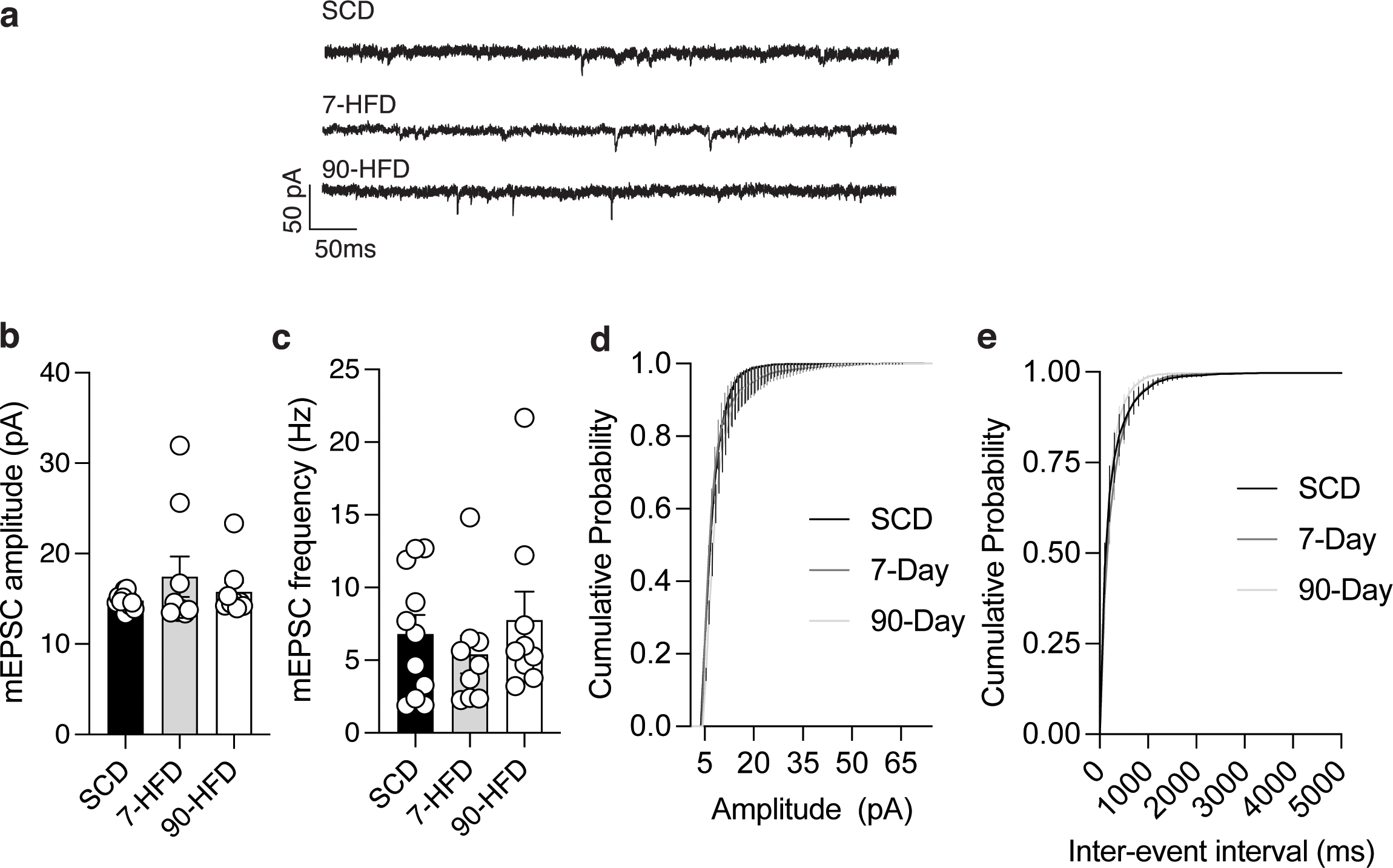
Short or long-term high fat diet exposure does not alter glutamatergic transmission. **a)** Representative recordings of mEPSCs in lOFC pyramidal neurons in SCD, 7-day HFD and 90-day HFD. **b)** There is no difference in the amplitude of mEPSCs between SCD (n/N=11/4), 7-day HFD (n/N=9/4) and 90-day HFD (n/N=9/3). One-way ANOVA: F (2, 26) = 1.02, P=0.37. **c)** There is no difference in the frequency of mEPSCs between SCD (n/N=11/4), 7-day HFD (n/N=9/4) and 90-day HFD (n/N=9/3). One-way ANOVA: F (2, 26) = 0.57, P=0.57. **d)** Cumulative probability plots for amplitude of mEPSCs between SCD (n/N=11/4), 7-day HFD (n/N=9/4) and 90-day HFD (n/N=9/3). There was a significant difference in cumulative probability of amplitude between groups: Kruskal-Wallis test, P = 0.022. **e)** Cumulative probability plots for inter-event interval of mEPSCs between SCD (n/N=11/4), 7-day HFD (n/N=9/4) and 90-day HFD (n/N=9/3). There was no significant difference in cumulative probability of inter-event interval between groups: Kruskal-Wallis test, P = 0.085.

We next tested if pyramidal neuron hyperexcitability is due to changes in GABAergic transmission. We measured lOFC pyramidal neuron excitability in the presence of picrotoxin, a GABA_A_ receptor antagonist. In picrotoxin, there were no differences in the excitability of pyramidal neurons between mice on SCD, and 7- or 90-day HFD (Figure 4a-c). In contrast to that observed in Figures 2g,h and j, AP width, AHP width and rheobase was not changed in the presence of picrotoxin in 90-day HFD-fed mice (Figure 4g,h,j). Furthermore, there was an increase in input resistance between SCD and 90-HFD (Figure 4k), but there was no difference in other membrane properties between groups (Figure 4d-j). These data suggest that increased pyramidal firing in mice exposed to short- and long-term HFD is modulated by inhibitory GABA_A_ receptors and may be due to decreased inhibition.

**Figure 4:**
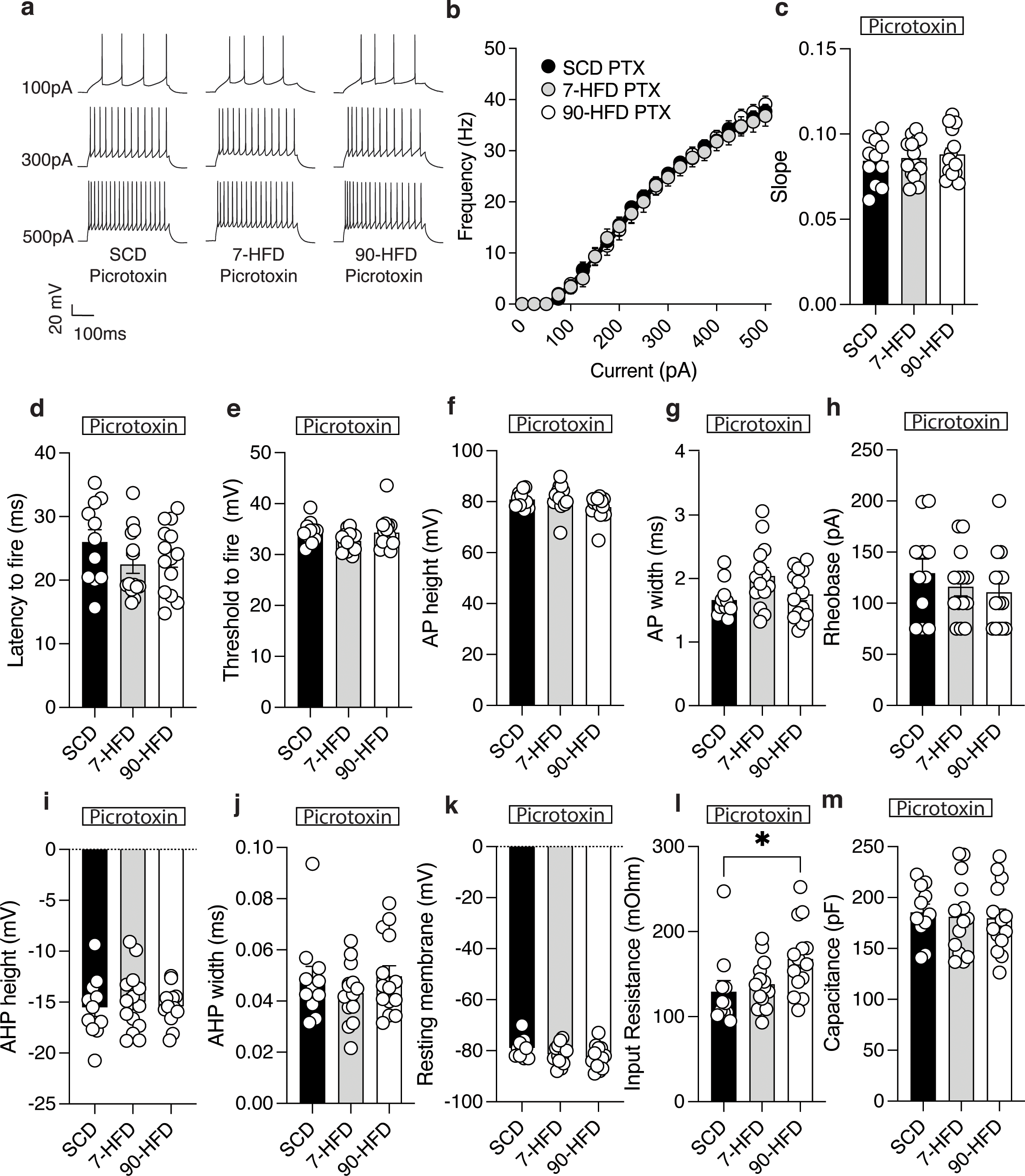
In the presence of a GABA_A_ receptor antagonist, picrotoxin, there are no differences in high-fat diet induced excitability. **a)** Representative recordings of action potentials (AP) in the presence of picrotoxin [100μm] observed at 100pA, 300pA and 500pA current steps from lOFC pyramidal neurons of standard control diet, 7-day HFD, or 90-day HFD. **b)** In the presence of picrotoxin, diet exposure: SCD (n/N=11/5), 7-day (n/N=14/3) or 90-day (n/N=14/3) HFD did not alter excitability as indicated by frequency-current (F-I) frequency of action potentials (AP) at current injections from 0 pA to 500pA. Two-way ANOVA: Diet effect F (2, 36) = 0.055, P=0.95, pA injected effect: F (2.70, 97.24) = 740.8, P<0.0001 ****, Diet x pA injected interaction: F (40, 720) = 0.46, P=0.99. **c)** In the presence of picrotoxin, diet exposure: SCD (n/N=11/5), 7-day (n/N=14/3) or 90-day (n/N=14/3) HFD did not alter slopes derived from a non-linear regression fit. One-way ANOVA: F (2, 36) = 0.25, P=0.78. **d)** In the presence of picrotoxin, diet exposure: SCD (n/N=11/5), 7-day (n/N=14/3) or 90-day (n/N=14/3) HFD did not alter the latency to fire. One-way ANOVA: F (2, 36) = 1.27, P=0.29. **e)** In the presence of picrotoxin, diet exposure: SCD (n/N=11/5), 7-day (n/N=14/3) or 90-day (n/N=14/3) HFD did not alter the threshold to fire. One-way ANOVA: F (2, 36) = 2.02, P=0.15. **f)** In the presence of picrotoxin, diet exposure: SCD (n/N=11/5), 7-day (n/N=14/3) or 90-day (n/N=14/3) HFD did not alter the AP height. One-way ANOVA: F (2, 36) = 2.56, P=0.090. **g)** In the presence of picrotoxin, diet exposure: SCD (n/N=11/5), 7-day (n/N=14/3) or 90-day (n/N=14/3) HFD did not alter the AP width. One-way ANOVA: F (2, 36) = 3.16, P=0.055. **h)** In the presence of picrotoxin, diet exposure: SCD (n/N=11/5), 7-day (n/N=14/3) or 90-day (n/N=14/3) HFD did not alter the rheobase. One-way ANOVA: F (2, 36) = 0.75, P=0.48. **i)** In the presence of picrotoxin, diet exposure: SCD (n/N=11/5), 7-day (n/N=14/3) or 90-day (n/N=14/3) HFD did not alter the AHP height. One-way ANOVA: F (2, 36) = 0.17, P=0.85. **j)** In the presence of picrotoxin, diet exposure: SCD (n/N=11/5), 7-day (n/N=14/3) or 90-day (n/N=14/3) HFD did not alter the AHP width. One-way ANOVA: F (2, 36) = 0.7971, P=0.46. **k)** In the presence of picrotoxin, diet exposure: SCD (n/N=11/5), 7-day (n/N=14/3) or 90-day (n/N=14/3) HFD did not alter the resting membrane potential. One-way ANOVA: F (2, 36) = 2.51, P=0.095. **l)** In the presence of picrotoxin, input resistance in the 90-day HFD (n/N=14/3) was greater than SCD (n/N=11/5) or 7-day HFD (n/N=14/3). One-way ANOVA: F (2, 36) = 3.68, P=0.035*, Dunnett’s post hoc comparison showed a difference between SCD and 90-day HFD P= 0.030*.

### GABAergic synaptic transmission is decreased by long-term HFD exposure

To test if short- and long-term diet exposure influenced inhibitory synaptic transmission onto lOFC pyramidal neurons, we measured spontaneous inhibitory post synaptic currents (sIPSCs). Mice fed 90 days of a HFD had decreased frequency but not amplitude of sIPSCs (Figure 5a-c). This is consistent with our previous work recording mIPSCs in the lOFC after a 90D HFD in mice (Seabrook et al., 2023) or an obesogenic cafeteria diet in rats (Thompson et al., 2017; Lau et al., 2021). We also observed a change in cumulative probability of inter-event interval in the 90D HFD group compared to SCD or 7D HFD (Figure 5d). However, there was no change in sIPSC amplitude (Figure 5a-c), cumulative probability of amplitude (Figure 5e) or mIPSC frequency, cumulative probability of inter-event interval, amplitude, or cumulative probability of amplitude (Figure 5f-i) in lOFC from mice fed 7 days of a HFD, suggesting that decreased inhibitory synaptic transmission release probability may not be responsible for increased pyramidal neuron excitability in mice fed 7 days of HFD.

**Figure 5:**
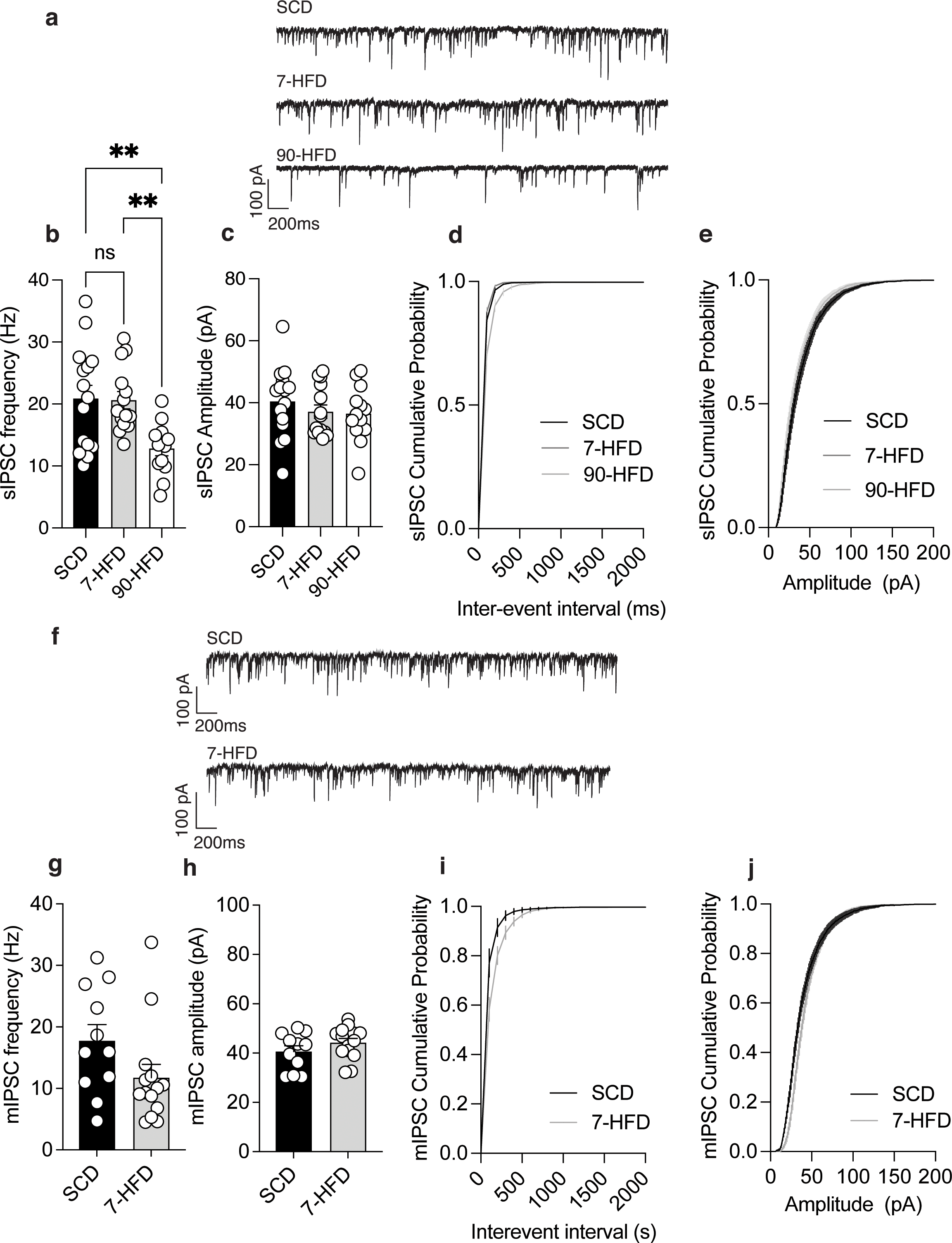
GABAergic synaptic transmission in the lOFC is decreased by long-term high fat diet exposure. **a)** Representative recordings of sIPSCs in lOFC pyramidal neurons in SCD, 7-day HFD and 90-day HFD. **b)** 90-day HFD (n/N=13/4) had decreased sIPSCs compared to SCD (n/N=15/5) and 7-day HFD (n/N=14/5). One way ANOVA: F (2, 39) = 7.17, P=0.0022. Tukey’s post hoc comparison test showed a significant difference between SCD and 90-day HFD P= 0.0045** and a significant difference between 7-day HFD and 90-day HFD p= 0.0071**. **c)** There is no difference in the amplitude of sIPSCs from SCD (n/N=15/5) 7-day HFD (n/N=14/5) and 90-day HFD (n/N=13/4). One-way ANOVA: F (2, 39) = 0.67, P=0.52. **d)** Cumulative probability plots for inter-event interval of sIPSCs SCD (n/N=15/5) 7-day HFD (n/N=14/5) and 90-day HFD (n/N=13/4). There was a significant difference in cumulative probability of inter-event interval between groups: Kruskal-Wallis test, P = 0.021*. **e)** Cumulative probability plots for amplitude of sIPSCs between SCD (n/N=15/5) 7-day HFD (n/N=14/5) and 90-day HFD (n/N=13/4). There was no significant difference in cumulative probability of amplitude between groups: Kruskal-Wallis test, P = 0.083. **f)** Representative recordings of mIPSCs recorded in TTX in lOFC pyramidal neurons in SCD or 7-day HFD. **g)** mIPSC frequency after 7-day HFD (n/N=14/5) was not significantly different from SCD (n/N=11/5). T-test: T (23) = 1.77, P=0.09. **h)** There is no difference in the amplitude of mIPSCs from SCD (n/N=11/5) and 7-day HFD (n/N=14/5). T-test: T (23) = 1.26, P=0.22. **i)** Cumulative probability plots for inter-event interval of mIPSCs SCD (n/N=11/5) and 7-day HFD (n/N=14/5). There was no significant difference in cumulative probability of inter-event interval between groups: Kolmogorov-Smirnov test, P = 0.75. **j)** Cumulative probability plots for amplitude of mIPSCs between SCD (n/N=11/5) and 7-day HFD (n/N=14/5). There was no significant difference in cumulative probability of amplitude between groups: Kolmogorov-Smirnov test, P = 0.05.

### Short- and long-term HFD exposure decreases tonic inhibitory transmission

Extrasynaptic GABA_A_ receptors tightly regulate neuronal firing in the cortex by providing persistent inhibition (Farrant and Nusser, 2005). To test if HFD exposure alters tonic inhibition, we measured the change in holding current of sIPSCs before and after gabazine, a GABA_A_ receptor antagonist, was applied to the slice. There was a decrease in gabazine-inhibited tonic current in mice fed both short- and long-term HFD (Figure 6a,b). In lOFC of mice fed 90 days but not 7 days of a HFD, there was a difference in the change of root mean square (RMS) noise, a proxy for tonically open GABA_A_ receptors (Bright and Smart, 2013) (Figure 6c). Taken together, these data suggest that both short- and long-term HFD exposure decreases tonic inhibitory drive onto lOFC neurons.

**Figure 6:**
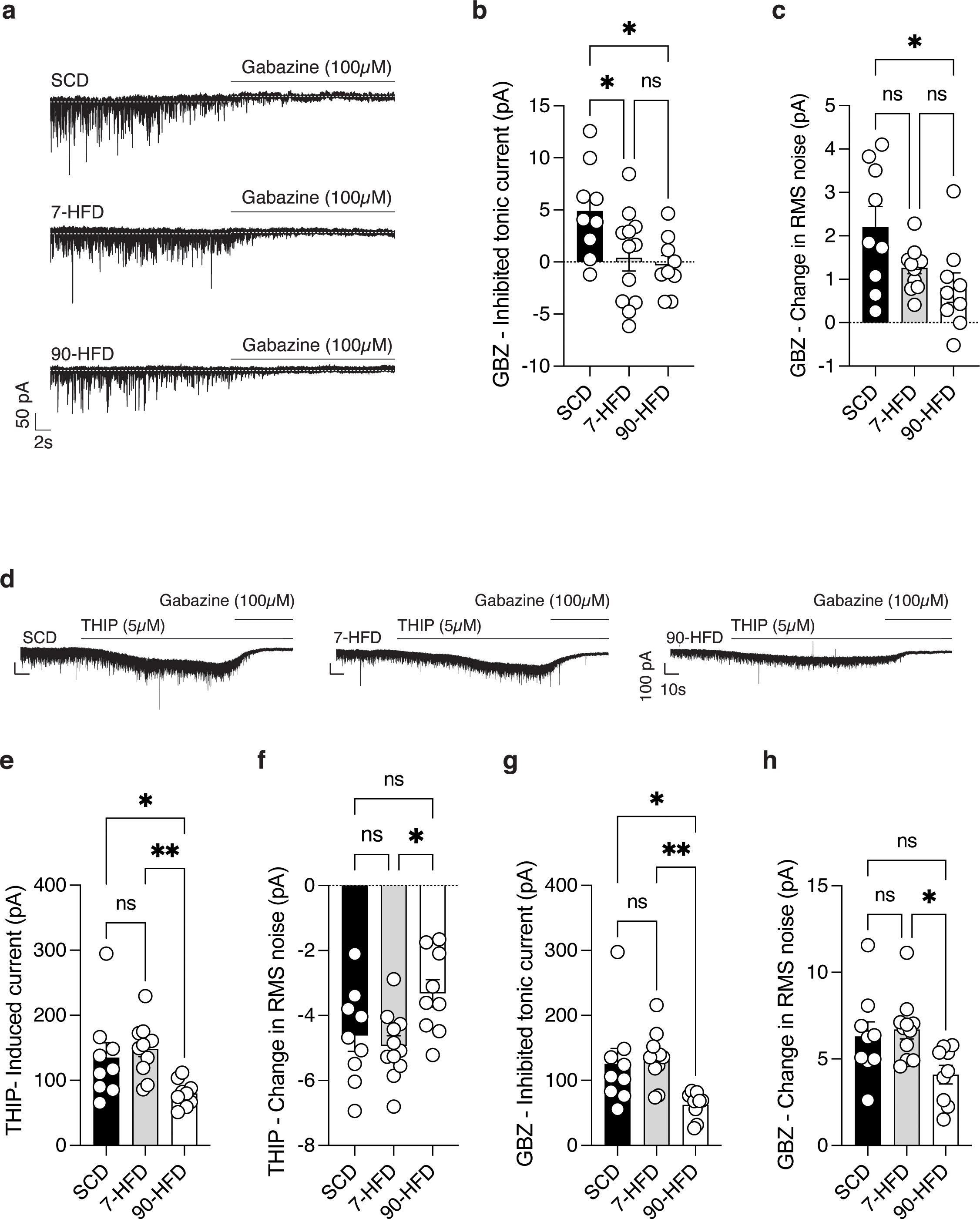
High fat diet exposure decreases tonic inhibitory transmission, but decreased 8 subunit containing GABA_A_ receptor currents in the lOFC only occurs with long-term HFD exposure. **a)** Representative recordings of tonic GABA in lOFC pyramidal neurons in SCD, 7-day HFD and 90-day HFD. **b)** Mice with 7-day (n/N=12/4) and 90-day (n/N=9/5) HFD exposure had decreased tonic inhibitory transmission compared to SCD (n/N=9/7). One-way ANOVA: F (2, 27) = 4.59, P=0.019 *, Tukey’s post hoc comparisons show a significant difference between SCD and 7-day HFD P= 0.045* and between SCD and 90-day HFD P= 0.027*. **c)** Mice with 90-day HFD (n/N=9/5) exposure had decreased tonic inhibitory transmission compared to SCD (n/N=9/7) and 7-day HFD (n/N=12/4). One-way ANOVA: F (2, 27) = 4.59, P=0.019*. Tukey’s post hoc comparisons show a significant difference between SCD and 90-HFD P=0.017*. **d)** Representative recordings of THIP-induced change in holding current in SCD, 7-day HFD and 90-day HFD lOFC pyramidal neurons. **e)** Mice with 90-day exposure to HFD (90-HFD n/N=9/3) have decreased delta subunit containing GABA_A_ receptor response to THIP compared to SCD (n/N=9/3) and 7-day HFD (n/N=11/5). One way ANOVA: F (2, 26) = 5.88, P=0.0078**. Tukey’s post hoc comparisons show a significant difference between SCD and 90-day HFD P= 0.045* and a significant difference between 7-day HFD and 90-day HFD P= 0.0077 **. **f)** Mice with 90-day exposure to HFD (90-HFD n/N=9/3) have a decreased THIP-induced change in RMS compared to 7-day HFD (n/N=11/5) but not SCD (n/N=9/3). One way ANOVA: F (2, 26) = 4.44, P=0.022*. Tukey’s post hoc comparisons show a significant difference between 7-HFD and 90-HFD P= 0.021*. **g)** Mice with 90-day HFD (n/N=9/3) exposure have decreased change in GBZ-induced tonic current compared to SCD (n/N=9/3) and 7-day HFD (n/N=11/5). One way ANOVA: F (2, 26) = 6.37, P=0.0056**. Tukey’s post hoc comparisons show a significant difference between SCD and 90-day HFD P= 0.026* and between 7-day HFD and 90-day HFD P= 0.0066**. **h)** Mice with to 90-day HFD (n/N=9/3) exposure have decreased change in RMS noise compared to SCD (n/N=9/3) and 7-day HFD (n/N=11/5). One way ANOVA: F (2, 26) = 4.69, P=0.018*. Tukey’s post hoc comparisons showed a significant difference between 7-day HFD and 90-day HFD P= 0.019*.

Extrasynaptic GABA_A_ receptors typically contain the delta subunit (Scimemi et al., 2005). As such, we tested the hypothesis that impairment of tonic conductance in 7- and 90-day HFD exposed mice was mediated by the delta subunit by testing the response to the delta subunit preferring agonist, gaboxadol (THIP). THIP-induced currents were decreased in 90-day but not 7-day HFD fed mice (Figure 6e,f). This was associated with a decrease in the change in RMS noise in the lOFC of mice fed a 90-day HFD (Figure 6f). There was also a decrease in gabazine inhibited tonic current in mice fed 90 days of a HFD as well as a decrease in gabazine induced change in RMS noise (Figure 6g,h). This suggests that the decrease in tonic current in 90-, but not 7-day HFD exposure, could at least in part be due to a decrease in delta subunit-containing GABA_A_ receptors.

### Increased excitability of pyramidal neurons, and decreased tonic inhibitory transmission is transient in short-but not long-term diet exposure

We next tested if the effects of short- and long-term diet exposure was transient or long lasting. We hypothesized that mice exposed to a 7-day HFD would have transient cellular adaptations, whereas 90-day HFD exposure leading to obesity and metabolic disfunction would have long-lasting effects. To test this, we replaced the HFD with the lower caloric density standard control diet for 7 days immediately after 7- and 90-days of HFD exposure (Figure 7a). 7 days after removal of the HFD, the long-term diet mice had increased excitability of lOFC pyramidal neurons as indicated by the F-I plot (Figure 7b-c) and excitability slope (Figure 7d) compared to 7-day HFD and SCD fed mice. Furthermore, consistent with action potential characteristics observed immediately after diet exposure, we found a significant decrease in rheobase and AHP width in the 90-day HFD group compared to SCD (Extended data Figure 7-1e,g). Notably, there was also a decrease in latency to fire and an increase in input resistance in the 90-day HFD group (Extended data Figure 7-1a,i). After removal of the HFD, there was no difference in tonic current revealed by gabazine between mice fed 7 days of HFD and standard control diet. However, 7 days after the 90-day HFD exposure there continued to be no tonic current revealed by gabazine (Figure 7e,f). There were no differences in gabazine-induced change in RMS noise between groups after removal of the HFD (Figure 7g). Taken together, these data demonstrate that lOFC pyramidal neurons are sensitive to HFD exposure, and changes in firing and tonic GABA observed after a short-term exposure are transient. In contrast, pyramidal neuron hyperexcitability and reduced tonic GABA adaptations in the lOFC occurring with long-term HFD exposure leading to metabolic dysfunction and weight gain are long lasting.

**Figure 7:**
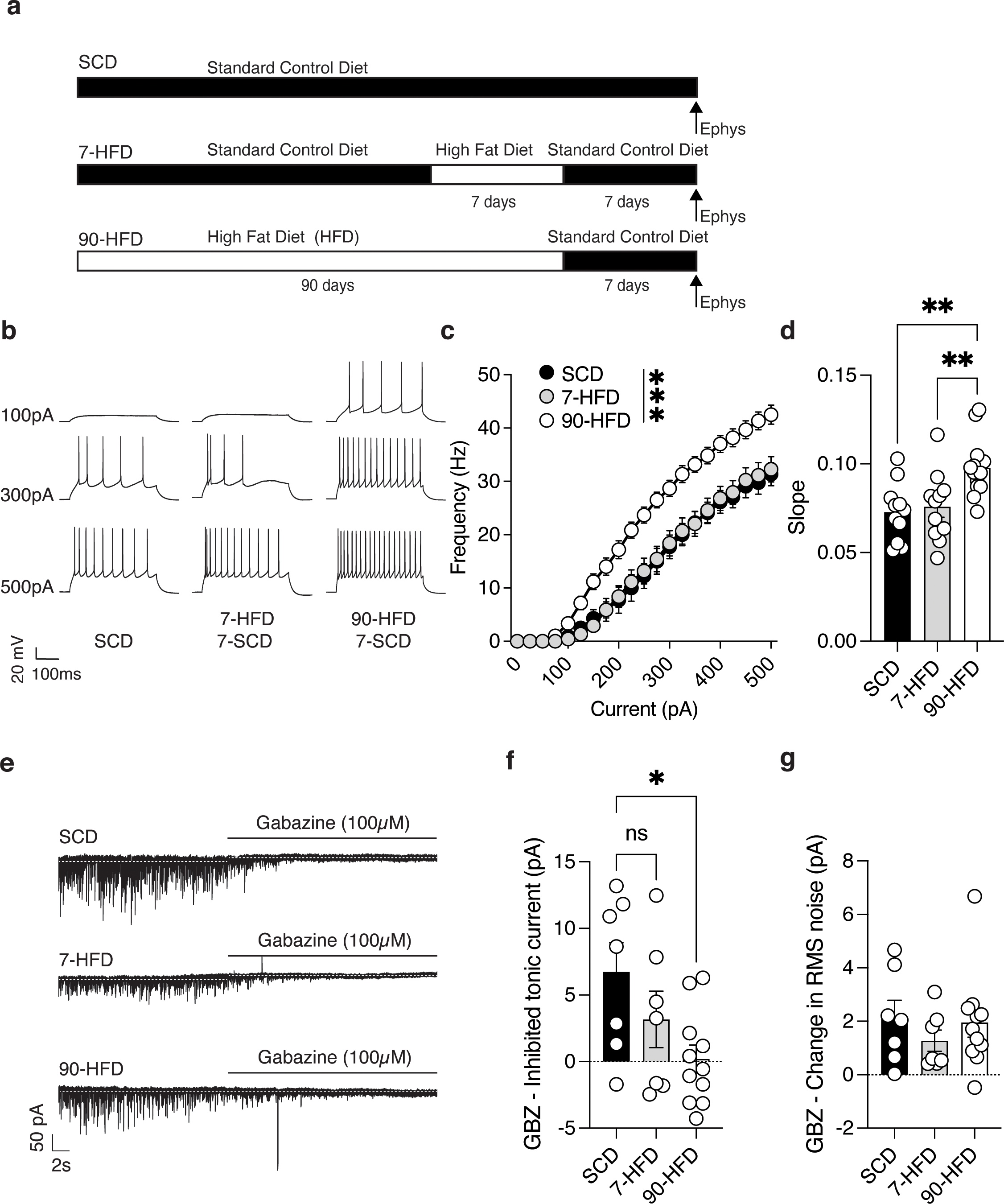
Increased excitability of pyramidal neurons, and decreased tonic inhibitory transmission is transient in short-but not long-term diet exposure. **a)** Schematic of high fat diet exposure and removal. **b)** Representative recordings of action potentials observed at 100pA, 300pA and 500pA current steps from lOFC pyramidal neurons in SCD, 7-day HFD and 90-day HFD after 7 days removal of HFD. **c)** After 7 days removal of HFD, 90-day HFD (n/N =15/3) had increased excitability of lOFC pyramidal neurons compared to SCD (n/N=11/5) and 7-day HFD (n/N =11/3) as indicated by frequency-current (F-I) at current injections from 0 pA to 500pA. Two-way ANOVA: Diet effect: F (2, 34) = 11.56, P=0.0001 ***, pA injected effect: F (2.13, 72.37) = 481.7, P<0.0001**** Diet x pA injected interaction: F (40, 680) = 6.67, P<0.0001***. Action potential characteristics are described in Figure 7-1. **d)** After 7 days removal of HFD, 90-day HFD (n/N =15/3) had increased excitability of lOFC pyramidal neurons compared to SCD (n/N=11/5) and 7-day HFD (n/N =11/3) indicated by a non-linear regression fit. One-way ANOVA: F (2, 34) = 8.77, P=0.0008 ***. Tukey’s post hoc comparisons showed a significant difference between SCD and 90-day HFD P= 0.0019 **, and 7-day HFD and 90-day HFD P= 0.0064**. **e)** Representative recordings of tonic GABA in lOFC pyramidal neurons of SCD, 7-day HFD and 90-day HFD mice after 7 days removal of HFD. **f)** After 7 days removal of a HFD, SCD (n/N=7/5) had increased gabazine inhibited tonic current compared to 90-day HFD (n/N=11/3) but not 7-day HFD (n/N=7/3). One-way ANOVA: F (2, 22) = 3.91, P=0.035 *, Tukey’s post hoc comparisons showed no significant difference between SCD and 7-day HFD P=0.37, but a significant difference between SCD and 90-day HFD P= 0.028*. **g)** After 7 days removal of a HFD there was no difference between SCD (n/N=7/5), 7-day HFD (n/N=7/3) and 90-day HFD (n/N=11/3) in the change of gabazine-induced RMS noise. One-way ANOVA: F (2, 22) = 0.57, P=0.58.

**Figure 7.1.**
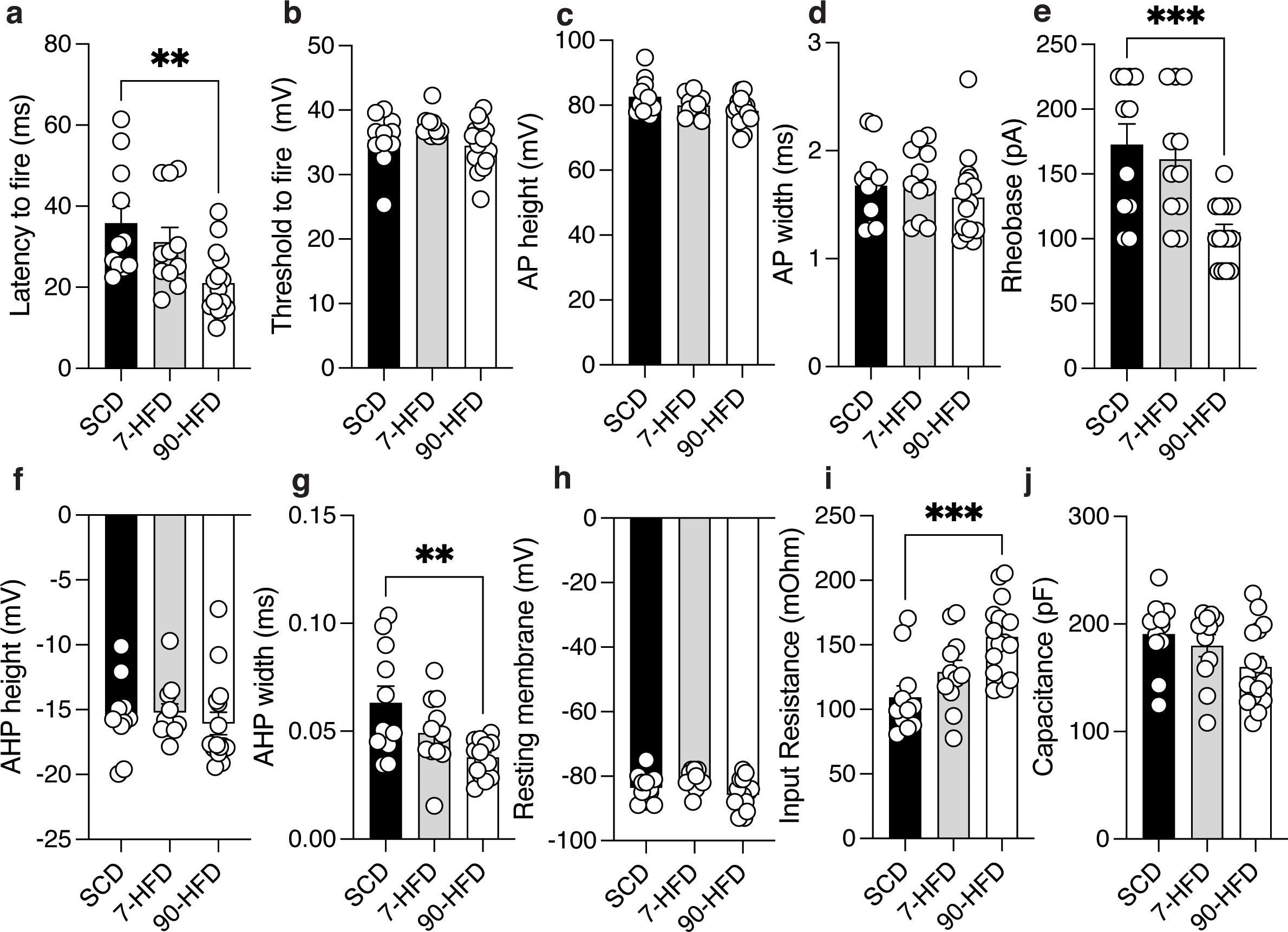
Short-term high fat diet exposure increases excitability of lOFC pyramidal neurons are transient compared to long-term high fat diet exposure. **a)** 90-day (n/N=15/3) HFD decreased the latency to fire compared to SCD (n/N=11/5) and 7-day (n/N=11/3) was not different from controls. One-way ANOVA: F (2, 34) = 609, P=0.0055. Dunnett’s multiple comparisons test: SCD vs 7-day HFD, P = 0.52, SCD vs 90-day HFD, P = 0.0038**. **b)** Diet exposure SCD (n/N=11/5), 7-day (n/N=11/3) and 90-day (n/N=15/3) HFD did not alter the threshold to fire. One-way ANOVA: F (2, 34) = 2.5, P=0.096. **c)** Diet exposure SCD (n/N=11/5), 7-day (n/N=11/3) and 90-day (n/N=15/3) HFD did not alter the AP height. One-way ANOVA: F (2, 34) = 2.99, P=0.06. **d)** Diet exposure SCD (n/N=11/5), 7-day (n/N=11/3) and 90-day (n/N=15/3) HFD did not alter the AP width. One-way ANOVA: F (2, 34) = 0.62, P=0.54. **e)** 90-day (n/N=15/3) or 7-day (n/N=11/3) HFD had decreased rheobase compared to SCD (n/N=11/5). One-way ANOVA: F (2, 34) = 10.14, P=0.0004. Dunnett’s post hoc comparison test showed a significant difference between SCD and 90-day HFD, P = 0.0005***, but no difference between SCD and 7-day HFD, P= 0.74. **f)** Diet exposure SCD (n/N=11/5), 7-day (n/N=11/3) and 90-day (n/N=15/3) HFD did not alter the AHP height. One-way ANOVA: F (2, 34) = 0.28, P=0.76. **g)** 90-day (n/N=15/3) HFD or 7-day (n/N=11/3) HFD had decreased AHP width compared to SCD (n/N=11/5). One-way ANOVA: F (2, 34) = 6.68, P=0.0036**. Dunnett’s post hoc comparison test showed a significant difference between SCD and 90-day HFD, P = 0.0017**, but no difference between SCD and 7-day HFD, P= 0.12. **h)** Diet exposure SCD (n/N=11/5), 7-day (n/N=11/3) and 90-day (n/N=15/3) HFD altered the resting membrane potential. One-way ANOVA: F (2, 34) = 3.93, P=0.029. However, a Dunnett’s post hoc comparison test showed no significant differences between SCD and 7-day HFD, P = 0.29 or SCD and 90-day HFD, P = 0.32. **i)** 90-day (n/N=15/3) HFD or 7-day (n/N=11/3) HFD had increased input resistance compared to SCD (n/N=11/5). One-way ANOVA: F (2, 34) = 8.28, P=0.0012**. Dunnett’s post hoc comparison test showed a significant difference between SCD and 90-day HFD, P = 0.0006***, but no difference between SCD and 7-day HFD, P= 0.21. **j)** Capacitance was similar between SCD (n/N=11/5), 7-day (n/N=11/3) and 90-day (n/N=15/3) HFD. One-way ANOVA: F (2, 34) = 2.54, P=0.09.

### Impairments in devaluation occur with short term diet exposure

Our previous work demonstrated that decreased GABAergic synaptic transmission led to an impairment in satiety or conditioned taste avoidance outcome devaluation and that boosting GABAergic tone in obese mice could restore performance on the outcome devaluation task <u>(Seabrook et al., 2023)</u>. Because we observed decreased tonic GABA after only 7 days of HFD exposure, we asked if there were differences in performance on the outcome devaluation tasks in these mice. Mice were conditioned over 3 sessions to associate either grape or orange flavoured gelatine with lithium chloride (LiCl)-induced malaise (Figure 8a). After 3 pairing sessions, mice consumed less of the LiCl-paired flavour. Mice were then fed 7 days of HFD or LFD and exposed to the unpaired (valued) and paired (devalued) flavours in a counterbalanced fashion. LFD fed mice demonstrated a conditioned taste avoidance to the LiCl-paired flavour, whereas 7-day HFD exposed mice consumed similar amounts of valued and devalued gelatine (Figure 8b). The strength of devaluation, represented by the revaluation index, was significantly different between LFD and 7-day HFD fed mice (Figure 8c) even though there was no difference in body weight (LFD: 32 ± 1g, 7-day HFD: 35 ± 1g, t(10)=0.52, P=0.61). To assess if the devaluation was due to a floor effect of gelatine consumption in the HFD group, we compared the total gelatine consumed in the valued and devalued states between groups. The total gelatine consumed was not different between groups (Figure 8d), suggesting that the devaluation was not due to a floor effect. We next tested how 7-day HFD exposed mice performed on the satiety-induced outcome devaluation task. Body weights were significantly different between SCD (30 ± 0.6, n = 8), 7-HFD (31 ± 0.7, n = 8) and 90-HFD mice (47 ± 0.7, n = 6). One way ANOVA, F(2,19) = 180.4, P < 0.0001. Tukey’s multiple comparison tests indicates significant differences between SCD and 90-HFD (P<0.0001) and 7-HFD and 90-HFD (P<0.0001), but not SCD and 7-HFD (p = 0.28), indicating again that 90-day HFD increases bodyweight from SCD controls, whereas 7-day HFD does not. Mice were trained to lever press for liquid sucrose on a random ratio (RR) 20 schedule of reinforcement, whereby sucrose deliver followed on average the 20^th^ lever press and then placed on a 7-day HFD or continued on SCD for the training and devaluation testing days (Figure 8f). 90-day HFD mice had diet exposure throughout training and testing. Mice were pre-fed sucrose (devalued state) or water (valued state) prior to the devaluation test (Figure 8e) where they expected delivery of sucrose solution in a non-reinforced test session. SCD-fed mice were sensitive to devaluation and responded less in the devalued condition compared the valued condition. In contrast, mice fed either 7- or 90-HFD had impaired outcome devaluation (Figure 8g), although this effect was considerably stronger in the obese mice (Figure 8g, h). Taken together, these data indicate that even a short-term exposure to HFD influences performance on outcome devaluation tasks.

**Figure 8.**
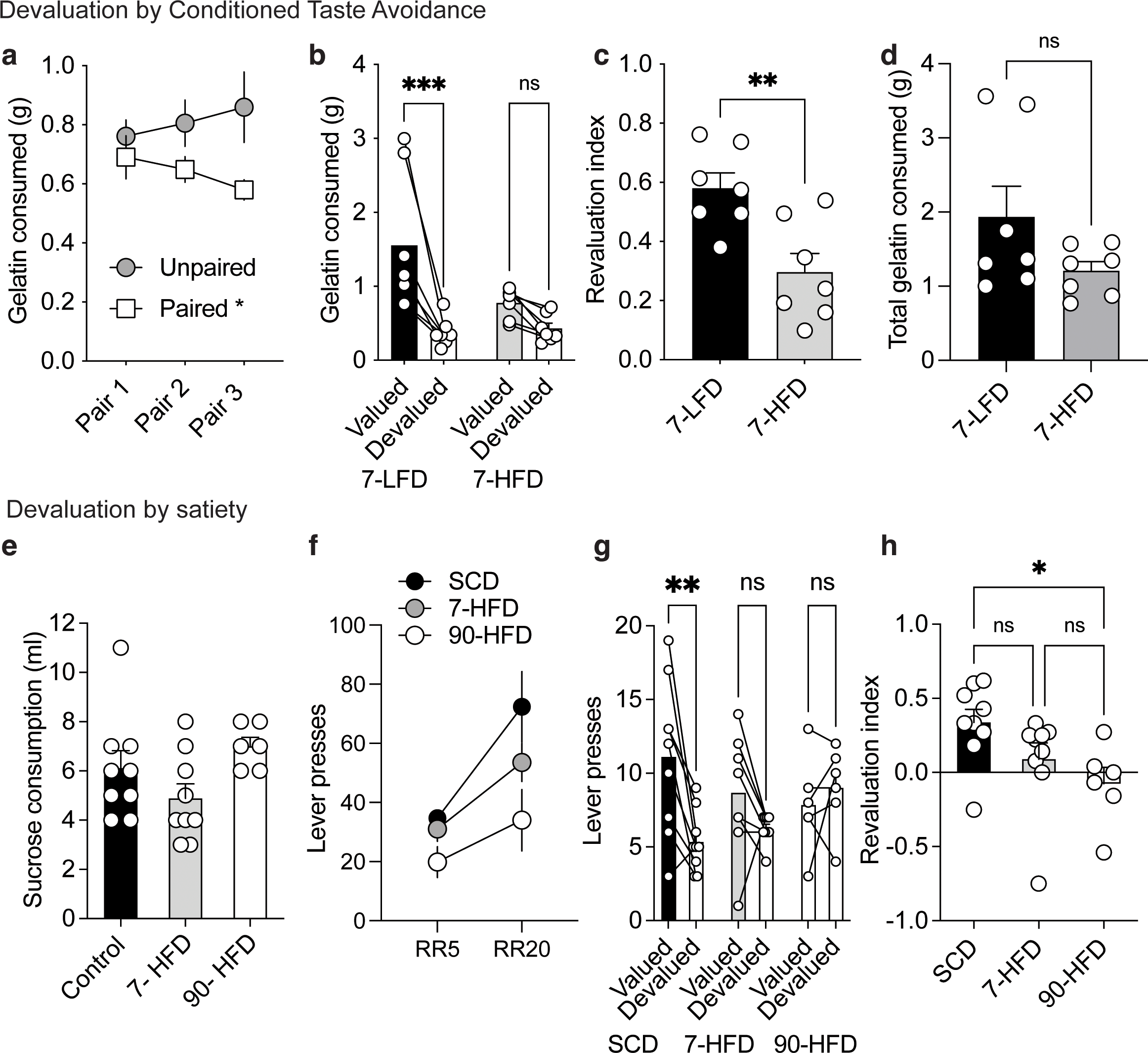
Exposure to a HFD induces deficits in outcome devaluation. **a)** Consumption (g) of the valued (unpaired) and devalued (paired gelatine during the three days of taste avoidance conditioning (pre-diet). RM Two-Way ANOVA: Day x devaluation interaction: F (2,48) = 1.79, p=0.18, Day effect: F (2,48) = 0.011, p=0.99, Valued vs. devalued effect: F (1, 24) = 4.94, p=0.036. Holm’s Sidak’s multiple comparisons test: Pairing Day 3: p = 0.018*. Data are presented as mean values +/-SEM. **b)** Consumption (g) of the valued and devalued gelatine during the CTA test after exposure to the diets (7-LFD n=7, 7-HFD n = 7). RM Two-way ANOVA: Diet x devaluation interaction: F (1,12) = 6.98, p = 0.02*. Devaluation effect: F (1, 12) = 23.26, p = 0.0004***, diet effect: F (1, 12) = 2.80, p=0.12, Holm’s Sidak’s multiple comparisons 7-LFD: p=0.0004***, 7-HFD: p=0.15. Bars represent mean values and symbols represent individual responses. **c)** Revaluation index of 7-LFD (n=7) and 7-HFD (n=7) mice. Unpaired t-test: t _(12)_ =3.45, p=0.0048. Data are presented as mean values +/-SEM and symbols represent individual responses. **d)** Total consumption (g) of the valued and devalued gelatine during the CTA 7-LFD (n = 7) and 7-HFD (n= 7) mice. Unpaired t-test: t _(12)_ =1.67, p=0.12. Data are presented as mean values +/-SEM and symbols represent individual responses. **e)** SCD (n=9), 7-HFD (n = 9) and 90-HFD mice (n=6) pre-test sucrose consumption was not different between groups. One-way ANOVA: F (2, 21) = 2.69, p=0.09. **f)** Training schedule on RR5 and RR20 for SCD (n=9), 7-HFD (n=9) and 90-HFD (n=6). RM two-way ANOVA: Ratio effect, F (1, 21) = 19.11. P=0.0003 ***, Diet effect F (2, 21) = 4.763, P=0.0197 *, Ratio x diet interaction F (2, 21) = 1.459, P=0.2552. Data are presented as mean values +/-SEM. **g)** SCD (n=9) mice displayed devaluation as indicated by decreased lever presses in the devalued compared to the valued condition. 7-HFD (n=9) or 90-HFD (n = 6) did not display devaluation as indicated by comparable lever presses in the valued and devalued conditions. RM two-way ANOVA: Devaluation effect, F (1, 21) = 7.54, p=0.012*, Diet effect F (2, 21) = 0.31, p=0.74, Diet x devaluation interaction, F (2, 21) = 5.16, p=0.015*, Holm-Sidak’s multiple comparisons test, SCD: p= 0.0012**, 7-HFD: p = 0.17, 90-HFD: p = 0.49. Data are presented as mean values (bars) and symbols represent individual responses. **h)** The revaluation index ((lever presses valued – lever presses devalued)/ (lever presses valued + lever presses devalued)) is significantly different between SCD (n=9), 7-HFD (n=9) and 90-HFD (n = 6). One way ANOVA, F(2,21) = 3.80, P = 0.039*. A Tukey’s multiple comparisons test indicates a significant difference between SCD and 90-HFD (p = 0.036). Data are presented as mean values +/-SEM with individual responses overlaid.

## Discussion

Chronic consumption of calorie dense foods can lead to weight gain, metabolic dysfunction, and concurrent health disorders. Lean and obese individuals overeat energy dense foods high in fats and sugar despite satiety signals or known health implications. Obesity alters synaptic function in the lOFC (Thompson et al., 2017; Lau et al., 2021; Seabrook et al., 2023) and impairs satiety-induced devaluation, an effect that is restored by increasing GABAergic tone in the lOFC of obese mice (Seabrook et al., 2023). We found that both short- and long-term HFD exposure increases excitability of lOFC pyramidal neurons with distinct inhibitory differences. With short-term HFD exposure, this effect was associated with a decrease in tonic inhibition, as glutamatergic and GABAergic release probability was not altered. Furthermore, while hyperexcitability and decreased tonic inhibition was transient after short-term diet exposure, these effects persisted only in the long-term diet exposure after returning mice to the standard control diet. The lasting hyperexcitability of pyramidal neurons in the lOFC of mice with long-term HFD exposure could be due to decreased synaptic and tonic inhibitory tone, consistent with previous reports (Seabrook et al., 2023). Finally, even after a short-term HFD exposure, there were impairments in outcome devaluation consistent with early onset changes in hyperexcitability. Here, we show that short-term HFD exposure does not alter body weight, and transiently increases the excitability of and decreases the tonic inhibition of lOFC neurons. Notably, populations of OFC neurons respond to sucrose rewards (Schoenbaum et al., 1998), scale with hunger (de Araujo et al., 2006) and the pleasantness of the food reward (Tremblay and Schultz, 1999). Taken together, these results point to a potential mechanism underlying how initial exposure to a HFD could contribute to further non-homeostatic eating.

### HFD exposure increases pyramidal neuron excitability in the lOFC

Although both short-and long-term HFD exposure increased excitability of lOFC pyramidal neurons compared to a standard control diet, this effect was graded such that neurons from mice with long-term diet exposure had an even greater excitability slope than the short-term exposure. The effects of obesity on lOFC neuronal firing are consistent with a previous report in obese mice fed a HFD (Seabrook et al., 2023). Furthermore, that these obesity-induced changes in lOFC firing are blocked by inhibitory synaptic transmission blockers is also consistent with that in mice (Seabrook et al., 2023) and rats with extended access to a cafeteria diet (Thompson et al., 2017). Notably, we observed significant changes in AP width and AHP width with diet exposure that are typically governed by potassium conductance. It is possible that this is due to an increase in depolarization state which would reduce the number of potassium channels open. While we did not observe this reflected in the resting membrane potential, consistent with our previous work (Seabrook et al., 2023), we did see a decrease in rheobase, which is also reflective of an increase in excitability state. These effects were blocked by picrotoxin and the increase in AHP width and rheobase were still present 7 days after removal of HFD only in the 90-day HFD group. We also observed an increase in input resistance in the 90-day HFD group both in the picrotoxin group as well as 7 days after 90-day HFD exposure, suggesting additional intrinsic mechanisms may also be at play. We did not observe a change in mEPSCs onto lOFC neurons of 7-day or 90-day HFD exposed mice. This is in contrast to that observed in rats fed extended access to a cafeteria diet, where mEPSC frequency onto lOFC pyramidal neurons was reduced due to increased extrasynaptic glutamate action at presynaptic metabotropic glutamate receptors 2/3 (Lau et al., 2021). It is possible that there are species or diet differences in the effects of diet on excitatory inputs. Alternatively, others have observed two populations of lOFC pyramidal neurons that differ based on electrophysiological characteristics in rats (Badanich et al., 2013). Thus, is possible that one of these populations may be overrepresented in rats compared to mice or vice versa and we were biased to different populations of pyramidal neurons in the lOFC of mice compared to rats. Regardless, changes in mEPSC do not necessarily reflect any changes in network activity and are unlikely to contribute to increased firing of lOFC neuron after diet exposure.

Short-term exposure to calorically dense food can alter synaptic transmission and neuronal activity in several brain regions. For example, one day exposure to sweetened high fat food induces synaptic strengthening of excitatory inputs onto ventral tegmental area dopamine neurons that can last at least a week (Liu et al., 2016). A one week exposure to a palatable high fat western diet produces an endocannabinoid-dependent short-term depression of excitatory synapses onto orexin neurons (Linehan et al., 2018), and increased excitatory synaptic transmission onto melanin-concentrating hormone neurons of the lateral hypothalamus (Linehan et al., 2020). The effects of short-term HFD exposure were transient in the lateral hypothalamus (Linehan et al., 2018, 2020). In the nucleus accumbens, a 10 day exposure to a junk food diet increases expression of calcium permeable AMPA receptors and has opposing effects on cellular excitability of medium spiney neurons from obesity susceptible compared to obesity resistant rats (Oginsky et al., 2016; Oginsky and Ferrario, 2019). While synaptic transmission does not appear to be affected by diet exposure in the motor or visual cortices (Lau et al., 2021), 7 days of a HFD impaired LTP field potentials in the prefrontal cortex (Shrivastava et al., 2021). It is possible that later access to the short-term HFD at P143-P150 compared to initiation of diet exposure during early adulthood (P60-P150) could impact the differential effects observed in electrophysiological characteristics. It is well established that early life and juvenile diet exposure have differential effect on brain and behaviour when compared to adult diet exposure (Boitard et al., 2012, 2014; Noble and Kanoski, 2016), which is why we chose only the adult period to initiate short and long term diet exposure. However, less is known how early adulthood vs mid adulthood influences diet-induced changes in the brain.

### HFD exposure alters GABAergic signalling

Several lines of evidence suggests that HFD exposure duration differentially alters GABAergic signaling. Application of the GABA_A_ antagonist, picrotoxin, reduced the increased excitability, including AP width, AHP width and rheobase observed in 7-day or 90-day HFD exposed mice. Further, there was decreased tonic GABA onto pyramidal neurons compared to standard chow fed rats. Because tonic GABA signalling can strongly modulate neuronal activity (Farrant and Nusser, 2005), a diet-induced reduction in tonic GABA may be a mechanism driving the increase in pyramidal neuron excitability. Tonic GABA is mediated by extrasynaptic GABA_A_ receptors typically containing α5 and delta subunits (Farrant and Nusser, 2005). We noted a strong response to the delta-containing GABA_A_ receptor agonist, THIP, in the lOFC of standard chow and 7-day HFD exposed mice, suggesting that there is significant expression of delta-containing GABA_A_ receptors in the lOFC. However, this effect was diminished in the lOFC of 90-day HFD exposed mice, indicating a reduced number or function of delta-containing GABA_A_ receptors in obese mice. Additionally, this suggests that mechanisms influencing tonic GABA function are different with short-term and long-term diet exposure. For example, the reduction in tonic GABA after 7-days of HFD could be due to reduced activity at delta-subunit containing receptors with no change in delta-subunit expression and prolonged diet exposure could lead to a reduction in delta-subunit expression. The response to the selective delta-containing GABA_A_ agonist THIP was not different between SCM and 7-day HFD, however a maximal dose was used which may override small changes in the efficacy of delta-containing GABA_A_ receptor function. Alternative hypotheses underlying reduced tonic GABA after 7 day exposure could be changes in α5 GABA_A_ subunit-containing receptor number or function, reduced reuptake of extrasynaptic GABA, or decreased perineuronal nets, as observed in rats exposed to a HFD (Dingess et al., 2018) leading to reduced firing of GABAergic neurons (Balmer, 2016) and disinhibited pyramidal neurons (Slaker et al., 2015). Future experiments will be aimed at testing these ideas.

In contrast to tonic GABAergic signaling, phasic signalling occurs when GABA_A_ receptors, typically containing α1, α2, α3 and/or ψ subunits, are located within the synapse and mediate point-to-point GABAergic transmission typically measured by inhibitory post synaptic currents (Farrant and Nusser, 2005). We did not observe changes in the frequency and amplitude of spontaneous or miniature IPSCs in the lOFC after 7 days of HFD. While we did not measure the mIPSCs after 90 days HFD in this study, we found that sIPSC frequency was reduced after long-term HFD exposure, consistent with a reduction in GABA release probability reported previously in obese mice (Seabrook et al., 2023) and rats (Thompson et al., 2017; Lau et al., 2021). Taken together, short-term and long-term diet exposure differentially influences GABAergic function in the lOFC, whereby tonic inhibition is decreased in both durations of diet exposure, but only mediated by reduced delta-containing GABA_A_ receptor function in obese mice and synaptic GABAergic transmission is only reduced after long-term HFD exposure.

Additional mechanisms underlying a shift in GABAergic function with diet induced obesity include regulation of parvalbumin (PV)-containing interneurons either directly or through changes to the perineuronal net (PNN) surrounding these neurons. The infralimbic prefrontal cortex and the ventral orbitofrontal cortex have reduced perineuronal net (PNN) staining surrounding parvalbumin (PV) expressing interneurons after 21 days of a HFD (Dingess et al., 2018). PNNs tightly regulate PV interneuron excitability and a reduction in PNN is associated with a decrease in PV interneuron excitability and has been reported in the medial prefrontal cortex (Slaker et al., 2015). Thus, it is possible a decrease in PV interneuron excitability may underlie reduced GABA release probability or tonic GABA observed with HFD diet exposure that could lead to disinhibition of pyramidal neurons. Furthermore, rats fed 8 weeks of a HFD decreased GABA levels in the frontal cortex (Sandoval-Salazar et al., 2016). Interestingly, there were no differences in PNN intensity or number in female rats exposed to a HFD, suggesting sex differences on diet-induced effects in the lOFC that should be explored in future studies. PV expression in prelimbic but not infralimbic PFC interneurons is also influenced by exposure to a sweetened HFD during the adolescent period in rats (Reichelt et al., 2015, 2019; Baker and Reichelt, 2016). This could be due to a decrease in the number of PV expressing neurons, although exposure to a cafeteria diet in rats did not influence the number of PV expressing neurons (Thompson et al., 2017). Alternatively, this may be due to diminished expression of the parvalbumin calcium binding protein, an effect that could lead to a decrease in the firing rate of PV-containing GABAergic neurons. Future experiments could address these questions.

Our previous work demonstrated that decreased in GABAergic input to lOFC pyramidal neurons leading to disinhibition of pyramidal neurons induced artificially with inhibitory DREADDs in lean mice or by diet induced obesity, can then lead to an impairment in outcome devaluation. Here, we replicated the effect that impaired satiety-induced devaluation occurred in 90-day HFD mice. However, we show that this effect occurs earlier on in HFD exposure. After 7-day HFD, both sickness-induced devaluation and satiety-induced devaluation were impaired. Moreover, this effect was graded, such that the strength of outcome devaluation by selective satiety was greater in the 90-day HFD exposure compared to 7-day HFD exposure. We also noted that the degree of disinhibition of pyramidal neurons was also greater in the 90-day HFD exposure compared to 7-day HFD exposure, suggesting that the cellular changes observed here may underlie the behavioural performance on the devaluation task. Although, we have not demonstrated that these changes are causally related in this study.

### Limitations

One caveat to our study is that the changing from chow to a HFD diet may induce a stress response. Our procedure aims to reduce neophobia by providing a small sample of HFD 48h prior to the diet switch. Furthermore, we do not see a drop in body weight with 7 days of a HFD, which one might expect with a stressor. Secondly, we used a low-fat diet (LFD) instead of SCD for the conditioned taste aversion experiment. Because LFD is less palatable than the SCD, it is possible that mice experienced a ‘worse than expected’ outcome that contributed to the recalled devaluation associated with the LiCl-paired flavour. However, mice did not lose weight with 7-day low fat diet exposure, suggesting that it was not a significant stressor. Thirdly, we did not test if mice were sated from the pre-test sucrose exposure prior to the satiety-induced devaluation task. Several other studies have identified that 0.5 - 2h sucrose exposure is sufficient to induce satiety for devaluation tasks (Johnson et al., 2009; Gourley et al., 2010; Lichtenberg et al., 2017). Consistent with this, we measured post-devaluation sucrose consumption in our previous study and found that mice drank less of the devalued flavour compared to the valued flavour, suggesting mice were sated during the pretest consumption period (Seabrook et al., 2023). An additional limitation of our study is for the last 7 days of the diet exposure, mice were socially isolated to avoid fighting that occurred with a staggered diet-delivery design. Thus, it is possible that effects of social isolation could contribute to our results. Notably, dorsal raphe dopamine neurons have increased AMPA/NMDA ratio when moved from group housed to isolated housing, yet social-isolation induced plasticity does not occur in VTA dopamine neurons (Matthews et al., 2016). In the OFC, 5 weeks adolescent social isolation from weaning increases the AMPA/NMDA ratio through postsynaptic mechanisms at lOFC-BLA synapses and decreases parvalbumin expression in the OFC of adolescent female, but not male mice (Kuniishi et al., 2022; Jeon et al., 2023). It is not known if these changes in OFC function occur with 7-day isolation during adulthood. However, several studies having indicated that adult brains are more resilient to social isolation compared to adolescents (Makinodan et al., 2012; Hinton et al., 2019; Rivera-Irizarry et al., 2020). Furthermore, we observe similar effects of 90 days HFD on firing and GABAergic synaptic transmission of lOFC neurons in this study, where mice were isolated for 7 days prior to recordings, compared to our previous study where all mice were group housed (Seabrook et al., 2023), suggesting that it is unlikely 7 days social isolation in adult male mice is influencing our electrophysiological responses. Finally, these experiments were performed in male mice. In obesity prone C57BL/6 mice, male mice are more susceptible to diet induced obesity with females taking a longer time course to become obese(Daly et al., 2022). Estrogen level(Bennett et al., 1998), sex-specific leptin resistance(Harris et al., 2003), and differences in gross locomotor activity(Benz et al., 2012) may contribute to sex differences in obesity development. Future experiments will explore how diet induced obesity impacts the lOFC of female mice.

### Conclusions

Our findings begin to elucidate the cellular mechanism on how lOFC pyramidal neurons are altered after HFD exposure. We observed that there are unique cellular adaptations in the lOFC with diet-induced obesity compared to short-term diet exposure. Understanding how acute and chronic consumption of calorically dense foods alters cortical regions has important implication for the self-regulation of food intake. Our previous work demonstrated that reduced GABAergic function in the lOFC of obese mice led to an impairment in the ability to use satiety to update the current value of the food reward (Seabrook et al., 2023). We replicate these effects and additionally show that this impairment occurs earlier on in diet exposure, suggesting that these behavioural changes are established prior to the development of obesity, tracking shifts in GABAergic function and disinhibition of pyramidal neurons in the lOFC. In conclusion, understanding the time course by which an energy dense diet alters neuronal activity and synaptic function in the OFC may reveal new mechanisms in the etiology of overconsumption and the development of diet-induced obesity.

## Acknowledgements

The authors would like to acknowledge the Hotchkiss Brain Institute advanced microscopy facility. This research was performed at the University of Calgary which is located on the unceded traditional territories of the people of the Treaty 7 region in Southern Alberta, which includes the Blackfoot Confederacy (including the Siksika, Piikuni, Kainai First Nations), the Tsuut’ina, and the Stoney Nakoda (including the Chiniki, Bearspaw, and Goodstoney First Nations). The City of Calgary is also home to Metis Nation of Alberta, Region III.

## Funding

This work is supported by a Koopmans Research Award, Mathison Centre for Research and Education Neural Circuits research grant, Tier 1 Canada Research Chair (950-232211) and Canada Institutes for Health Research Foundation Grant (CIHR FDN 148473 SLB). LTS was supported by a Harley Hotchkiss Doctoral Scholarship in Neuroscience and the Alberta Graduate Excellence Doctoral Scholarship. Colleen Peterson is supported by Alberta Innovates Graduate Studentship in Health Innovation.

## Author Contributions

LTS and CP performed electrophysiological experiments. LTS and CP fed mice. LTS, TK, AKJ, MS, and TT performed behavioural experiments. LTS, SL, and MA analyzed the data. LTS and SLB wrote first drafts of the manuscript. LTS, CP, DN, MA, MS, and SLB revised the manuscript.

## Conflict of Interest Statement

The authors declare no competing financial or other conflicts of interest.

## Notes

### Competing Interest Statement

The authors have declared no competing interest.

### Summary of Updates

New analyses added, updated figures

